# Let’s Get Physical: On Observing Force Perturbations of Proliferating Glioblastomal Cells

**DOI:** 10.1101/2023.01.24.525397

**Authors:** Jong Seto, Yong Chen

## Abstract

The growth and proliferation of mutant astrocyte cells are widely known characteristics in high grade glioblastoma multiforme (GBM) neurological cancers. The infiltrative tumor processes during glioblastoma development follow predefined routes and patterns dictated by biochemical and physiological environments in the brain. Specifically, at the cellular-level the glioblastoma-related astrocytes cells typically are occluded along neural fissures, neurophysiological interfaces as well as along available blood vessel routes for sources of nutrients to drive overall tumor tissue growth and development. Additionally, each individual astrocyte cell maintains a predisposition to aggregate into a supracellular aggregate with organioid-like properties. Here in this work, we attempt to understand the physical processes involved at the single cell level in forming these pre-tumor states and investigate the environmental forces that may perturb the formation of intermediate cellular assemblies in order to ultimately perturb overall tumor growth and development processes.

## Introduction

Many human disorders and disease, whether neurological or endocrine in origin, are found to be a result of the basis of subcellular and cellular aggregation processes. This includes Alzheimer’s with the aggregation of the beta-amyloid and tau protein to form plaques and tangles, Parkinson’s with the protein aggregation to create Lewy bodies intra-cellularly, as well as cataract related blindness with the nonspecific aggregation of the alpha crystalline protein subunits on the lens and symptoms of diabetes where high amounts of blood sugar causes systemic metabolic cell dysfunction. In several cases, the main dysfunction is related to an inability of a key protein to fold or regulate properly activity, subsequently, leading to inhibition of cellular activities or in some cases, leading to aggregation, inhibition of normal cellular growth and proliferation and, eventually cellular death in normal tissue.

As found with all tumors, the aggregation propensity at the cellular length-scale leads to the development and growth of cancers. In the case of glioblastoma multiforme (GBM), the most common human neurological cancer, is mainly driven by mutant astrocyte cells which undergo rapid growth and infiltration; invading into proximal neurological spaces normally occupied by other cells and tissues. Growth and infiltration widely acknowledged as following fissures and blood vessels. These masses often proliferate along white matter tracts, also known as Scherrer tracks, in the brain tissue and infiltrate into surrounding tissues [1–3]. The ability to infiltrate freely through extraneous spaces and occupied tissue by way of a pathophysiochemical toolbox inherent in the astrocyte cells. Specifically, these cells release active enzymes and molecules which act as matrix metalloproteases or inactivators of metalloprotease inhibitors such that the degradation of the neighboring extracellular matrices is accomplished to support the proliferative behavior of the nascent glioblastoma. Several groups have identified molecular and physical markers through magnetic resonance imaging and histopathological maps in attempts to resolve neuroanatomical areas that are highly prevalent to glioblastomas [4, 5]. Recently, specific isolated genetic mutations in the MGMT and PRMT5 genes in glioblastma-related astrocyte cells have been correlated to the histomorphologies found in the glioblastoma pathology [6, 7] [8], however, treatment of these glioblastomas still remain to be confined to treatment regimes pertaining to only radiotherapy and chemotherapy at the tissue [9] [10]. In this work, we demonstrate that intermediate formation states of glioblastoma can be observed by physical perturbations at the cellular level. This reveals a possible new area of study that can be utilized for halting tumor progression and growth.

## Results

Current cell culture methods have been optimized to reflect the need to accurately simulate the microenvironments that are found physiologically [11]. Through the use of custom made polymethylmethacrylate cell culturing wells fashioned with varying diameters for culturing of glioblastoma-related astrocyte cells, the surface tension on the substrate of the cultured cells can be modulated to observe surface tension effects on the growth of astrocyte cells. With the assumption that the liquid-surface interface is homogenous and only the radii of curvature are different, the Young-Laplace Equation can be used to quantify the contribution of the surface tension in each well. Specifically, the surface tension can be expressed as a function of Laplace pressure:

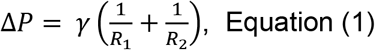

where P is the Laplace pressure, γ is the Surface Tension, R_1_ and R_2_ are the radii of the curvature. Since this is a liquid-surface interface, the Laplace pressures in varying diameters of the wells must be equal, resulting in the following relationships with the wells of varying diameters:

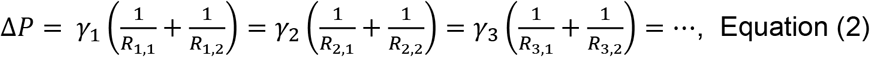

where R_1,1_, R_2,1_, R_3,1_,…,R_i,1_ are equal since only the diameter of the wells in one dimension change and can be expressed:

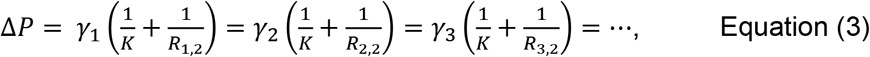

such that the surface tension is proportional to the radius of each well

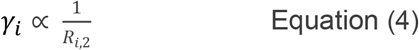

In our experiments, we performed glioblastoma-related astrocyte cell cultures in wells with diameters from 10 mm to 100 mm using identical volumes and cell numbers. The surface tension in each of these wells would scale accordingly as shown in Equation (4), such that by increasing the diameter of the wells from 10 mm to 100 mm, the surface tension would decrease proportionally. In fact, the surface tension in the 10 mm to the 100 mm well would be such that γ_100mm_ ~ 10 γ_10mm_. From our observations of culturing in wells with diameters of 10 mm, 50 mm, and 100 mm, we observe dramatic phenotypic differences in the growth of the astrocyte cells over the course of a 1 week cell culture. As shown in Figure 3 b, the cultures in the 100 mm diameter well shows less aggregation and formation of organoid-like structures compared to the 50 mm diameter well culture. In further comparisons, the 10 mm diameter well, with the smallest diameter and highest surface tension tested here, showed the largest single cellular mass of the other wells and the densest. From these observations, we preliminarily confirm that glioblastoma-related astrocyte cell growth and glioblastoma progression *in vitro* can indeed be affected by the physical environment in which these cells are initially seeded, revealing a possible route for new therapeutics.

## Discussion

A much less discussed pathobiology of glioblastomas is the hierarchical structuring schemes of the individual glioblastoma-related astrocyte cells that contribute to the formation of the invading tumor masses. At the single cell level, individual astrocyte cells demonstrate their ability to form cell-cell interactions readily, often forming interactions with multiple neighboring astrocyte cells. These interactions form the spheroid-like cellular structures observed in 48 hour cell cultures. Interestingly, these intermediate spheroid formations themselves display a kinetic equilibrium whereby these formations are approximately the same size and contain the same number of astrocyte cells. The homogeneity of these formations resemble the organoids which are derived from a variety of stem cell sources [12] [13]. These intermediate formations themselves then initiate a secondary growth stage where the surrounding edges of the “spheroid of astrocyte cells” spread into contact with other proximal “spheroids.” This can be observed over a course of a 1 week period in cell culture. Eventually, the spaces between these organoid-like structures begin to “fill-in” from fast cellular growth and an ever expansive cell spreading of astrocytes at the periphery of these structures. And the continued aggregation with surrounding organoid structures form the cell dense glioblastoma tissue masses detected by magnetic resonance imaging at the tissue level (Figures 1 and 2).

**Figure 1.**
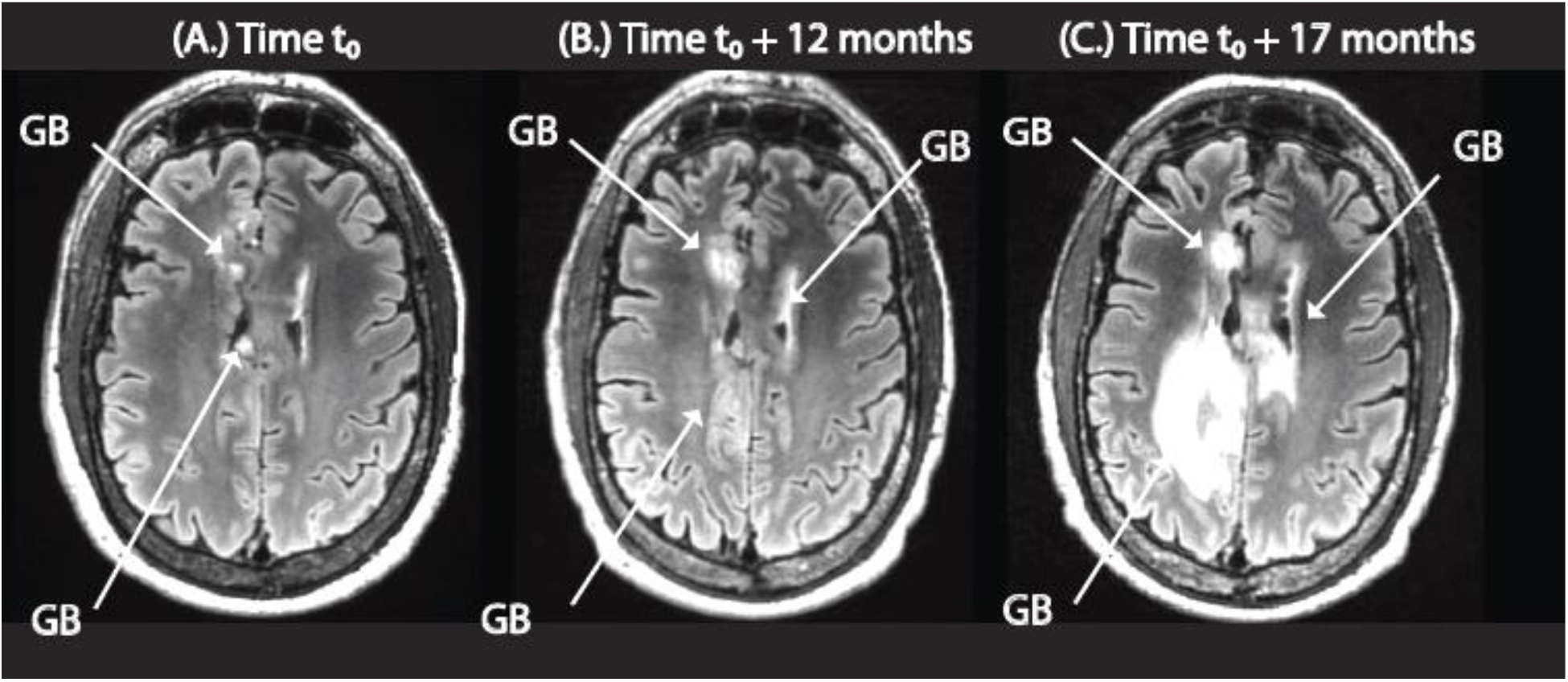
Magnetic Resonance Imaging of Glioblastoma Growth and Proliferation over time depicting high degrees of invasiveness and proliferation into healthy tissue (A.) Initial observation at time t_0_ (under T2 weighted imaging mode, fluid attenuated inversion recovery (FLAIR)) (B.) 12 months time point after t_0_ (C.) 17 months time point after t_0_

**Figure 2.**
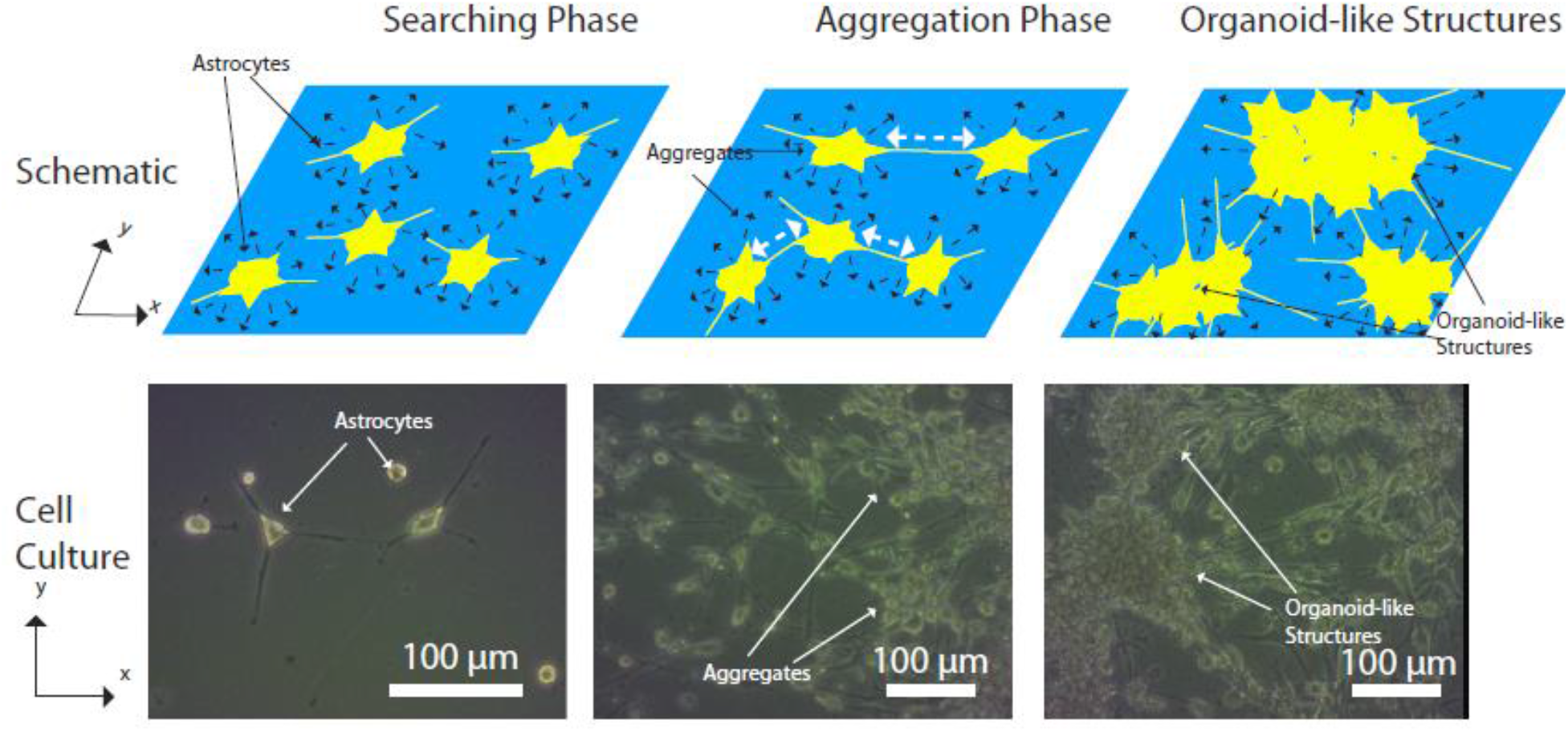
Growth of Glioblastoma – related astrocyte cells in cell culture and the progression to the diseased state as shown through schematic and optical images (A.) Astrocyte cells immediately after seeding (B.) Cells in culture after 48 hours (C.) Cells in culture after 1 week

**Figure 3.**
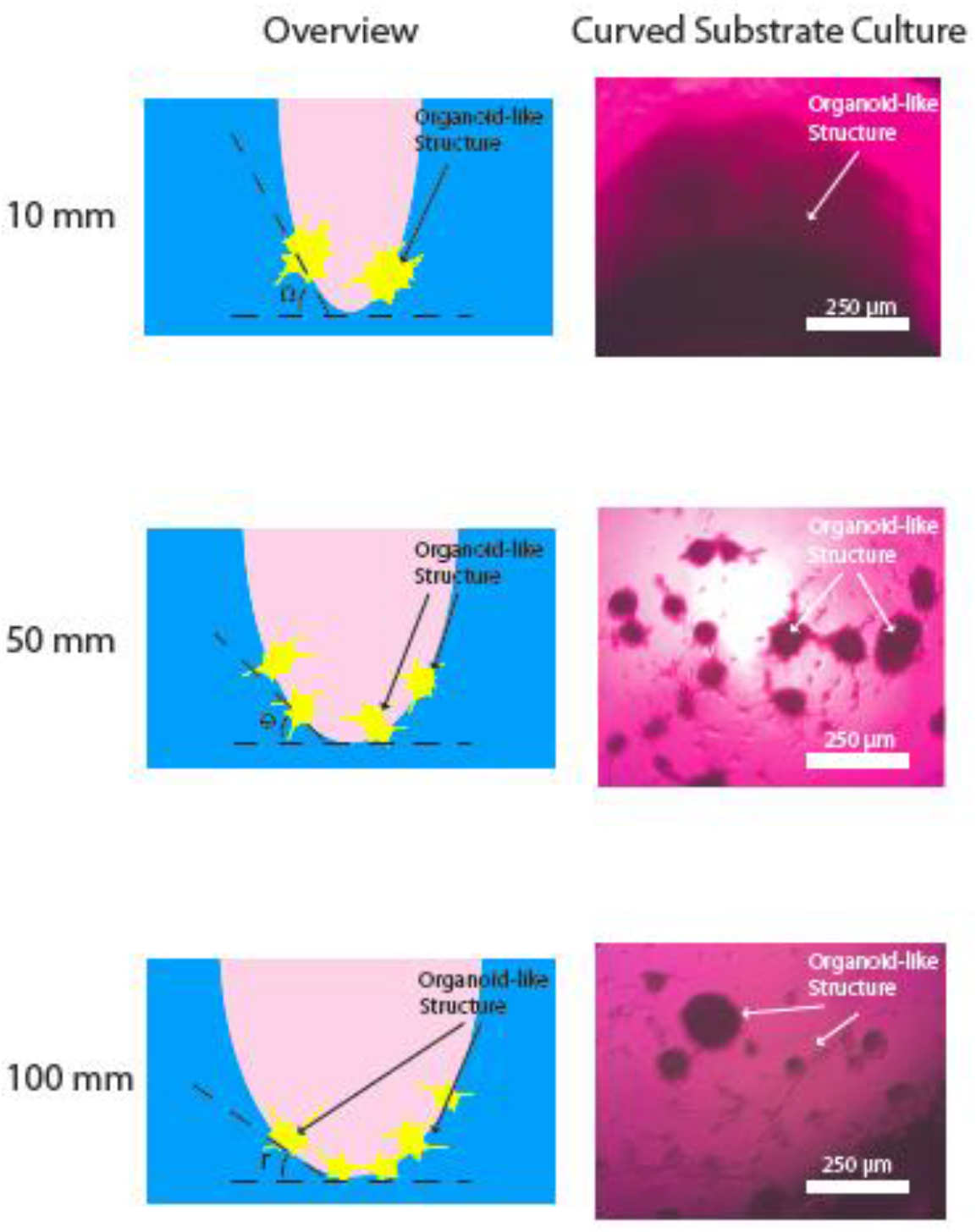
Surface tension effects on the growth and proliferation of glioblastoma – related astrocyte cells in culture such that Ω > θ > Γ.(A.) 10 mm (B.) 50 mm (C.) 100 mm

Using three-dimensional cell culturing techniques, normal and diseased states are simulated experimentally to recreate the physiologically-relevant scenarios of cellular organization found in tissues. Several groups are investigating the utility of these diverse methods of cell culturing to observe perturbed behaviors of normal cells with the intention to find aberrant structure or function at the single cell level [14] [15]. Here in this work, we observe how diseased astrocyte cells behave and organize with respect to a change of the physical environment at meso-scale (Figure 3).

We investigate the various length-scales of diseased cellular states here in order to understand how cellular organization can regulate diseased states at the tissue length-scale and propose options that address the fundamental underpinnings how diseased tissues persist and grow. Specifically, this preliminary work demonstrates that the mechano-sensory and transduction pathways within the astrocyte cells are involved in the formation of the organoid-like structures and may likely be connected to growth and development of the tumor tissue. Several groups have already implicated this system in normal cell group in various model systems to modulating a diversity of cellular function [16–18] [19–21]. And shown previously, scaffold geometry has a tremendous effect on new cellular growth patterns [22]. We confirm these single cell to tumor tissue transitions through these observations in perturbing the surface tension, aggregration ability of individual cells can be varied even in diseased states. By decreasing the surface tension on these glioblastoma related astrocyte cells, less single cells aggregate to form the large tumor tissue measured by MRI (Figure 3). In fact, by increasing the diameter from 10 mm to 50 mm, the surface tension decreases by approximately 20%. And further increasing the diameter from 50 mm to 100 mm, deceases the surface tension term approximately 40%. This results in the decreased cell aggregation ability and correspondingly, fewer organoid-like structures. In order to quantify this aggregation effect, a parameter ζ is used to measure the average size of the cell aggregates taking into account the size of these aggregates and the number of cellular aggregates. We find that ζ_10_ >> ζ_100_, further confirming our observations with these glioblastoma-related astrocytes in curved culture wells (Table 1).

**Table 1.** Aggregate sizes as a function of well diameter.

| Diameter of well (mm) | Average aggregate size, $\zeta$, (mm), | Standard Errors of Mean |
| --- | --- | --- |
| 10 | 1.46* | 0.033 |
| 50 | 0.072 | 0.00689 |
| 100 | 0.064 | 0.00833 |
| * Average aggregate sizes are statistically significant to other wells ($P = 0.012$) | | |

Further work to dissect the components of this mechano-sensitive response in glioblastoma-related astrocyte cells such as identifying cell surface markers that are intracellularly relevant in the diseased cell, sequencing of single cell genome and proteome to identify single nucleotide polymorphisms (SNPs) or displayed antigens, deciphering the signaling pathways which connect the mechano-sensitive response to tumorigenic states in these cells. Additionally, another aspect of great interest is the cellular interface of normal and diseased astrocyte cells and how these different populations coexist. Through utilizing updated methods of co-culturing of healthy and diseased cell states, interactions between the different cell populations can demonstrate how proliferation of diseased cells occur and through specific mechanisms [23]. This preliminary study attempts to investigate and correlate physically the molecular underpinnings that lead to these various chemical environments that cause cells to deviate from their normal states of activity.

## Materials and Methods

### Magnetic Resonance Imaging of Glioblastoma tissue

The Surbeck Advanced Imaging Laboratory at UCSF utilize standard clinical imaging protocols to image invading glioblastoma-related tissue in patients in accordance with UCSF IRB protocols. All images were acquired in a 3 Tesla MRI with 32 channel receiver system (GE Healthcare, Chicago, Illinois 60661, USA) and made anonymous in compliance with HIPAA standards and rules. MRI data acquired for this manuscript contains no identifiable personal traits or landmarks.

#### Construction of the curved substrates

A 80 × 50 × 20 mm (L × W × T) block of PMMA (Acrylic Plastic 24210-08, Electron Microscopy Sciences, Hatfield, PA 19440, USA) was cast and cured in a mold for 24 hours. Using a three-axis, custom built, computer numerical control (CNC) milling machine with various sized steel drill bits, diameters of 100 mm, 50 mm, 10 mm wells were drilled with a depth of 10 mm into the block of PMMA.

#### Glioblastoma cells cultures

Human glioblastoma cell line U87-MG was prepared in Dulbecco’s Modified Eagle Medium (DMEM) completed with 10% FBS and 1% penicillin/streptomycin at 37 °C with 5% CO_2_ supplementation for 3–4 days. After proliferated to confluence, cells were detached by trypsin at 37 °C for 3 min and centrifuged before re-suspended in a culture medium at a density of 5 × 10^6^ cells/mL.

#### Growth in curved substrates

An aliquot of 50 μL of harvested, resuspended cultured glioblastoma cells were placed into each curved substrate well making the number of cells in each well to be approximately 2.5 × 10^5^ cells/mL. An additional 50 μL of media was added to increase the final volume in each well to 100 μL. The wells were incubated in 37 °C for 48 hours before examination under the microscope. After 48 hours incubation, the glioblastoma-related organoid structures can be typically observed.

#### Optical Light Microscopy

An inverted optical light microscope (Zeiss Axiovert 200M, Carl Zeiss Microscopy GmbH, Carl-Zeiss-Promenade, Jena 07745, Germany) with a CCD (Evolution QEi Digital Camera, Media Cybernetics Inc., Silver Spring, MD 20910 USA) was used with 20X and 40X objectives to examine the cell cultures in the curved substrates. All captured images were analyzed with ImageJ (NIH, 9000 Rockville Pike, Bethesda, MD 20892, USA).

## Acknowledgements

Dr. Yadong Tang (ENS) is acknowledged for her technical advice and discussions on glioblastoma cell cultures. Marissa Lafontaine (UCSF), Adam Autry (UCSF), the Surbeck Laboratory for Advanced Imaging and the Quest sponsored Research Program in the Department of Radiology and Biomedical Imaging at UCSF are thanked for their aid in acquiring and analyzing magnetic resonance imaging of clinical glioblastoma patients. All clinical protocols followed UCSF IRB procedures.

## Author Contributions

JS conceived and designed experiments, performed experiments, analyzed data, and drafted the manuscript. YC also revised and drafted the manuscript.

SI Figure 1. T1 weighted imaging with gadolinium (A.) Initial observation at time t_0_ (B.) 12 months time point after t_0_ (C.) 17 months time point after t_0_

